# Decision-making shapes dynamic inter-areal communication within macaque ventral frontal cortex

**DOI:** 10.1101/2024.07.05.602229

**Authors:** Frederic M. Stoll, Peter H. Rudebeck

**Affiliations:** Nash Family Department of Neuroscience, Lipschultz Center for Cognitive Neuroscience and Friedman Brain Institute, Icahn School of Medicine at Mount Sinai, New York, NY, 10029, USA

**Keywords:** ventral frontal cortex, orbitofrontal cortex, agranular insula, reward, decision making, choices, outcome, functional connectivity

## Abstract

Macaque ventral frontal cortex is comprised of a set of anatomically heterogeneous and highly interconnected areas. Collectively these areas have been implicated in many higher-level affective and cognitive processes, most notably the adaptive control of decision-making. Despite this appreciation, little is known about how subdivisions of ventral frontal cortex dynamically interact with each other during decision-making. Here we assessed functional interactions between areas by analyzing the activity of thousands of single neurons recorded from eight anatomically defined subdivisions of ventral frontal cortex in macaques performing a visually guided two-choice probabilistic task for different fruit juices. We found that the onset of stimuli and reward delivery globally increased communication between all parts of ventral frontal cortex. Inter-areal communication was, however, temporally specific, occurred through unique activity subspaces between areas, and depended on the encoding of decision variables. In particular, areas 12l and 12o showed the highest connectivity with other areas while being more likely to receive information from other parts of ventral frontal cortex than to send it. This pattern of functional connectivity suggests a role for these two areas in integrating diverse sources of information during decision processes. Taken together, our work reveals the specific patterns of inter-areal communication between anatomically connected subdivisions of ventral frontal cortex that are dynamically engaged during decision-making.

## INTRODUCTION

The primate ventral frontal cortex plays a central role in guiding adaptive behavior during decision-making. When making a choice, neural activity within orbitofrontal cortex (OFC) and ventrolateral prefrontal cortex (vlPFC) represents the different attributes associated with the available options, such as the amount, effort, delay, risk, or probability that the option might be able to be obtained (Tremblay and Schultz, 1999; Padoa-Schioppa and Assad, 2006; Kennerley and Wallis, 2009; Kobayashi et al., 2010; Chau et al., 2015; Kaskan et al., 2017; Stoll and Rudebeck, 2024a). The OFC and vlPFC, are not, however, anatomically homogeneous areas and each encompasses a number of distinct subdivisions that have been defined on the basis of sulcal anatomy, cytoarchitecture, and receptor density (Walker, 1940; Barbas and Pandya, 1989; Morecraft et al., 1992; Carmichael and Price, 1994; Rapan et al., 2023). On top of this, anatomical tracing studies have revealed that each of these subdivisions receives a distinct set of projections from other parts of the brain (Barbas and Pandya, 1989; Carmichael and Price, 1995a, 1995b, 1996). Our previous neurophysiology recording study has also reported dissociable encoding patterns across the subdivisions of ventral frontal cortex (Stoll and Rudebeck, 2024a) and such differences in encoding appear to correspond to the effect of pharmacological inactivation of these subdivisions (Murray et al., 2015; Rudebeck et al., 2017). Altogether, this indicates specialization of function within distinct parts of the macaque ventral frontal cortex.

The anatomically distinct subdivisions of ventral frontal cortex are also densely interconnected with each other (Carmichael and Price, 1996). Based on this, as well as the patterns of connections coming from outside of the ventral frontal cortex, Carmichael and Price suggested that there might be specialized functional networks within ventral frontal cortex. Despite this, little is known about the patterns of functional communication between the subdivisions of ventral frontal cortex during decision-making. Prior work using resting-state fMRI in human and monkeys highlighted the existence of distinct networks within ventral frontal cortex based on their functional connectivity profiles with other regions (Kahnt et al., 2012; Kahnt and Tobler, 2017; Rapan et al., 2023). Given the role of ventral frontal cortex in adaptive behavior, resting-state connectivity might not provide an adequate way to understand the communication between areas during decision-making as functional interactions are likely shaped by the cognitive operations that are currently engaged (Zald et al., 2014). For example, a human neuroimaging study found that the connectivity between posterior OFC and ventromedial frontal cortex correlated with the sensory-specific changes in encoding induced by satiety (Howard and Kahnt, 2017). However, this human neuroimaging study and others like it are unable to discern the patterns of functional communication between anatomically defined subdivisions at the level of single neurons during decision-making due to the resolution of MRI.

To address this, we analyzed large-scale and high-density neurophysiological recordings that were made across eight distinct cytoarchitectonic areas of the macaque ventral frontal cortex, looking for time varying patterns of functional communication between areas. Recordings of single neuron activity were made while monkeys performed a two-alternative forced choice probabilistic task for different juice rewards. Taking this approach, we found specific patterns of functional communication between areas that were distinct from baseline periods. Notably, we found strong time-varying connectivity between areas 12o/12l and other subdivisions during stimulus and reward periods. Connectivity patterns were linked to the representation of decision variables and showed information flow in specific directions across the network of areas, with the more integrative areas, 12o and 12l, more likely to receive information from other subdivisions of ventral frontal cortex. Thus, our findings provide the first account of how subdivisions of ventral frontal cortex interact during decision making, highlighting specific roles for specific interareal functional communications.

## RESULTS

### Task, Behavior and Neural representation of decision variables

The behavior of the subjects has been reported in detail before (Stoll and Rudebeck, 2024a). In brief, we trained two monkeys to perform a two-choice probabilistic task in which each option was composed of a central gauge (more or less filled) signaling the probability at which a juice reward would be earned, and a colored frame indicating the juice flavor monkeys would receive if that option was selected (**Figure 1**). Monkeys reported their choice by fixating a response box located on each side of the screen. Following a feedback period in which both options were displayed again, a reward was delivered (or not) according to the monkeys’ choice (which defined the probability and flavor). Analyses of choices from 289 sessions (monkey M and X = 103 and 186 sessions) revealed a strong influence of the probability of receiving a reward on monkeys’ choice behavior (**Figure 1**). Outcome flavor also influenced monkeys’ choices, which was evident from the variable shift in the sigmoid functions from one session to another (see (Stoll and Rudebeck, 2024a) for detailed analyses on how flavor preference modulated choices and neuronal responses).

**Figure 1.**
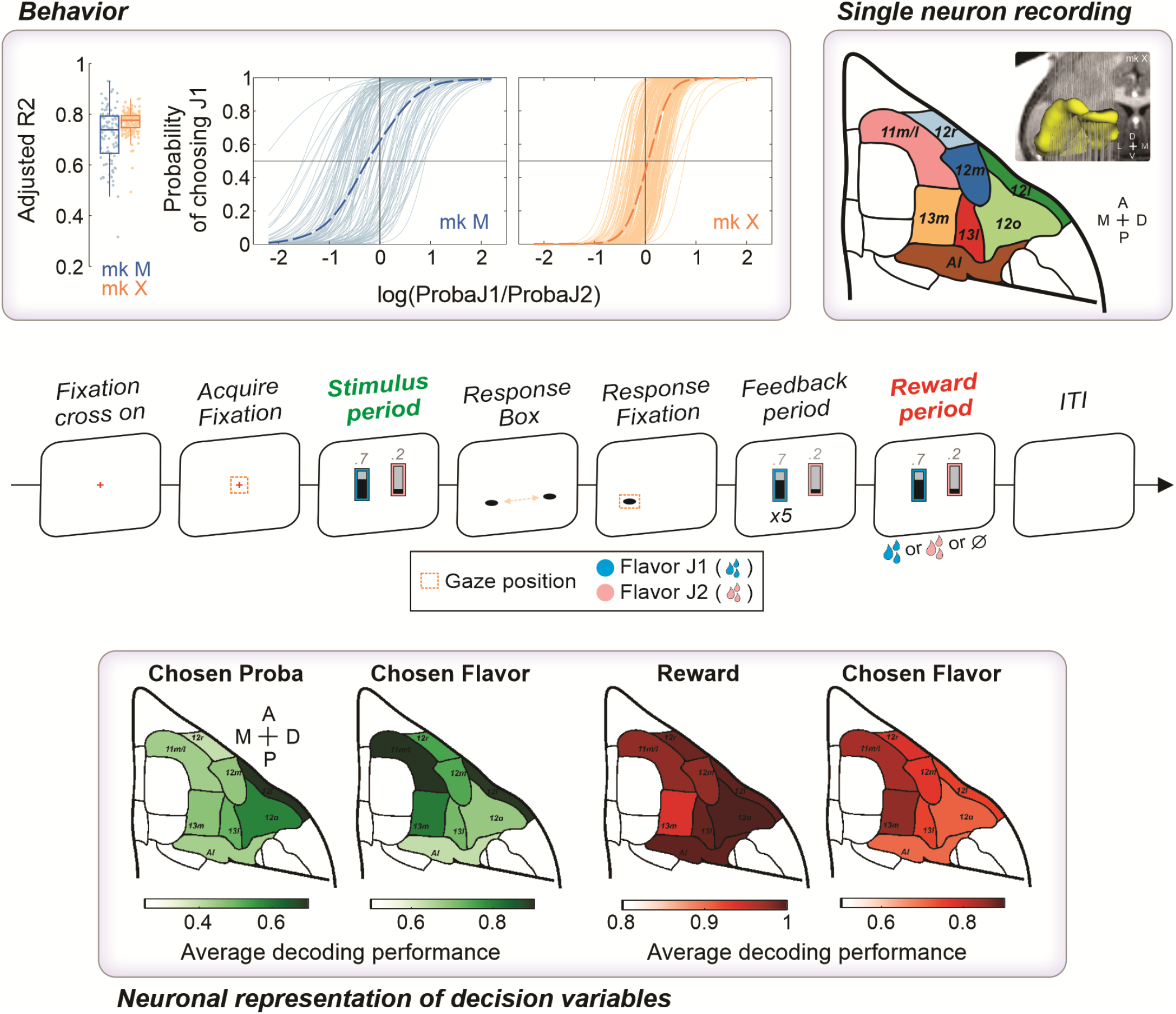
Overview of the task, behavior, recording and neuronal representations. Two monkeys performed a two-choice probabilistic task (center panel) in which they were asked to choose one of two possible stimuli, each comprised of a background color (indicating the outcome flavor that could be earned) and a central gauge (indicating the probability of receiving such outcome). Monkeys’ choices depended on the offered outcome probability (sigmoid function) and flavor (varying bias across sessions) (top left panel). We recorded the activity of 6,284 single neurons in 8 subdivisions of ventral frontal cortex (top right panel) while monkeys performed the task, revealing substantial variation in the representation of decision variables as assessed using pseudo-population decoding methods (bottom panel). During the stimulus period, 12l neurons more strongly represented both relevant decision variables (probability and flavor) compared to other areas, while 12o neurons were more specialized in the representation of chosen probability and 11m/l neurons in the representation of flavor. At the time of the reward delivery, most areas exhibited strong decoding of whether monkeys received or not the reward, while neurons in 11m/l and 13m strongly represented its flavor. Abbreviations: A=anterior, P=posterior, M=medial, L=lateral, D=dorsal, V=ventral, o=orbital, r=rostral. Adapted from (Stoll and Rudebeck, 2024b). See also **Table S1**.

While monkeys performed this task, we recorded the activity of 6,284 neurons across 8 cytoarchitectonic areas within ventral frontal cortex (**Figure 1** and **Table S1**, see (Stoll and Rudebeck, 2024b)). The precise recording locations were defined based on a combination of CT scans co-registered to MRIs and post-mortem immunohistology, following the parcellations previously reported by Price, Palomero-Gallagher, and colleagues (Carmichael and Price, 1994; Rapan et al., 2023). Our recordings covered subdivisions within vlPFC (areas 12r, 12r, 12l, 12o), OFC (areas 11m/l, 13m and 13l), and agranular insula (AI).

Previous work has repeatedly found that stimulus presentation and reward delivery influence the activity of single neurons in ventral frontal cortex (for instance, (Thorpe et al., 1983; Tremblay and Schultz, 1999; Padoa-Schioppa and Assad, 2006; Kennerley et al., 2009; Stoll and Rudebeck, 2024a)). Recently, we showed that representations are, however, not uniform across anatomically distinct subdivisions of ventral frontal cortex (Stoll and Rudebeck, 2024b). Specifically, neurons in 12l exhibited the strongest and most diverse representation of the parameters critical for the valuation of the different options during the task (**Figure 1**). Neurons in 11m/l more selectively encoded the chosen flavor whereas neurons in 12o more strongly represented the chosen probability of receiving a reward compared to other subdivisions. At the time of the reward delivery, all subdivisions represented whether a reward was delivered or not, with the strongest encoding found in area 12o. Neurons in area 11m/l and 13m were most likely to discriminate which reward flavor the subjects’ monkey expected or received, respectively. Here we looked at whether, and how, the different subdivisions communicate their representations with each other during the decision-making processes.

### Assessing functional connectivity within ventral frontal cortex

To assess inter-areal communication, we used cross-validated canonical component analysis (CCA), a method that allows the modelling of linear relationships between two multivariate sets of variables (Härdle and Simar, 2007; Semedo et al., 2020). Specifically, we used CCA to search for the linear combinations of activity of two sets of simultaneously recorded neurons from distinct areas so that the activity subspaces from these two areas were maximally correlated (see **STAR Methods**). We ensured our results were generalizable through cross-validation and focused on the strength of this correlation (canonical component correlation, referred to as *CC rho*) across every pair of areas in which we recorded at least 4 neurons simultaneously. Note that we focused on results observed in both monkeys and do not make strong statements regarding results from 12r and 12m as the recordings from these areas did not entirely overlap anatomically across monkeys (see number of sessions in **Table S2**).

During the baseline period, when monkeys held their gaze steady on the central fixation cross, we observed a relatively low and uniform zero-lagged connectivity across areas within ventral frontal cortex (**Figure 2A**). The highest observed connectivity during that time was between two neighboring areas, 12o and AI. The overall connectivity then increased as the trial progressed to the stimulus and reward periods. During the stimulus period, there was an increase in connectivity between area 12l and areas 11m/l, 13l and 12o, as well as between area 12o with areas 13l and AI (**Figure 2B-C**). As the trial progressed into the reward period, connections between 12o and areas 12l, 13l, and AI exhibited the strongest observed connectivity. Both monkeys showed strong connectivity between 12o and 12l, while the connectivity between 12o and 13l/AI was most prominent in monkey X (**Figure S1**). We also observed a dissociation over time, where neural activity within area 12l aligned more with area 13l than 12o during the stimulus period (and did so in a sustained way) while it aligned more with area 12o than 13l during the reward period (**Figure 2A-B**). Another clear pattern of connectivity was observed between area 12m and areas 12o and 11m/l, which was stronger in anticipation of the reward (feedback period) and slowly decreased after reward onset. Area 13m failed to show strong connectivity with any other subdivisions across the course of the trial, with the notable exception of an increased connectivity with area 11m/l, 13l, 12o and AI, peaking almost one second following the reward.

**Figure 2.**
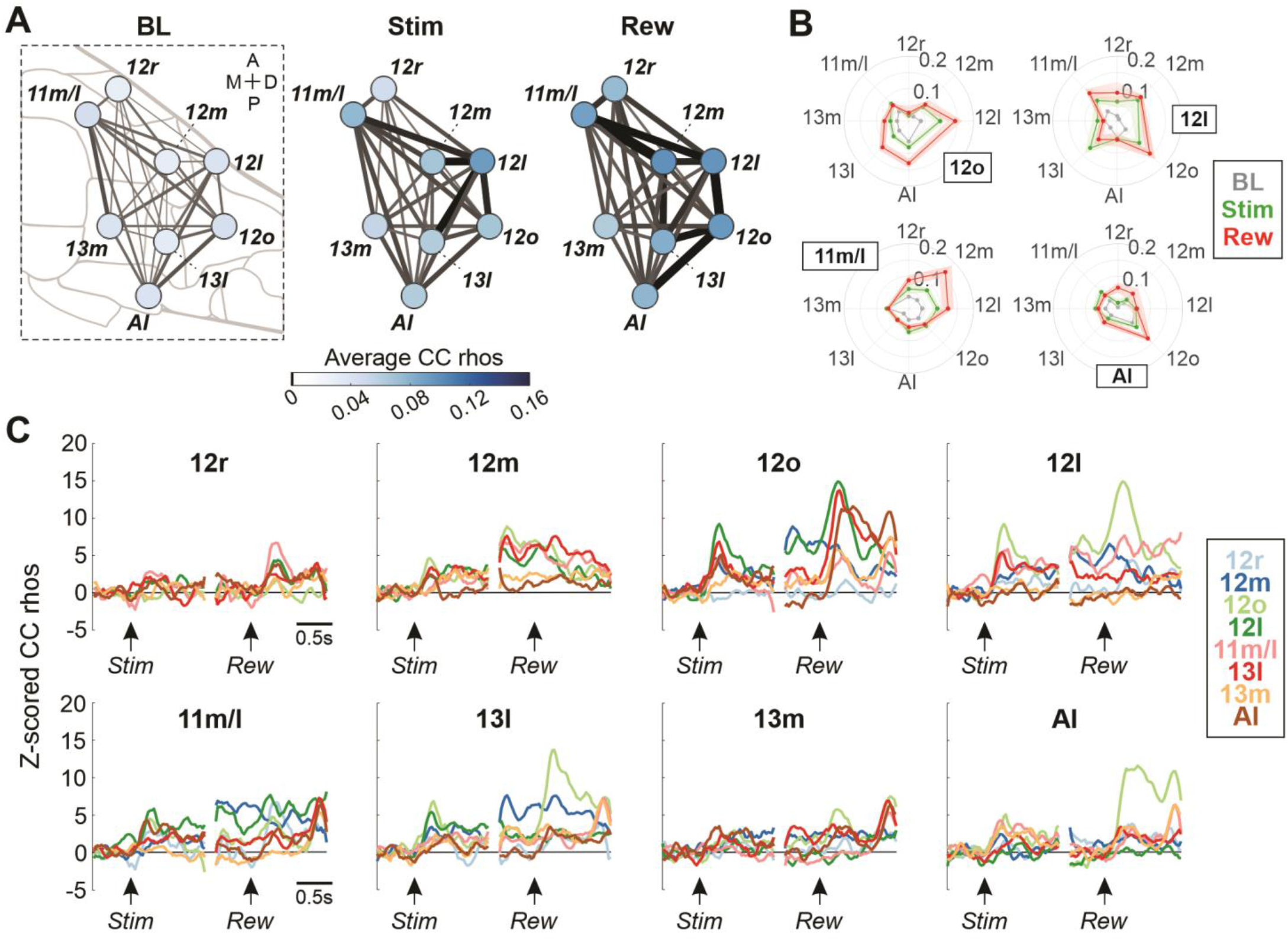
Functional connectivity using CCA. (**A**) Graph representation of the average zero-lagged correlation between canonical components extracted using cross-validated CCA for each pair of areas during the baseline, stimulus onset and reward onset periods (left to right), reported on a ventral view of the macaque frontal cortex. The thickness and darkness of each line represents the strength of the correlations (i.e., thicker and darker lines for stronger correlations). Each dot color represents the average correlation for a given area with all others. BL: baseline; Stim: stimulus; Rew: reward periods (**B**) Connectivity fingerprint for 4 example areas (12o, 12l, 11m/l and AI), showing the average zero-lagged correlation across sessions for the 3 considered time periods (baseline=grey, stimulus onset=green, reward onset=red). Shaded areas indicate s.e.m. (**C**) Time course of the average zero-lagged correlations (z-scored using the baseline period) around stimulus and reward onsets (arrows), for each area compared to all others. See also **Figure S1** and **Table S2**.

In summary, this pattern of results indicates that subdivisions of ventral frontal cortex do not communicate uniformly with one another during the task and instead exhibit specific and dynamic alignment of neural activity across subdivisions and time. In particular, vlPFC areas 12o and 12l exhibited strong time-varying connectivity with both anterior (11m/l) and posterior (13l and AI) subdivisions.

### Functional connectivity depends on task-related activity

Our finding that connectivity within ventral frontal cortex evolved over the course of the trial indicates that inter-areal communication is shaped by the cognitive processes that are currently being engaged. At the start of each trial, animals must first assess the attributes and value associated with each stimulus they can choose. As the trial progresses into the feedback and reward phase, they need to ensure that the outcome of their choices match their expectations and adjust their valuations accordingly. The specific attributes of the available options on each trial are encoded by neural activity within ventral frontal cortex (Thorpe et al., 1983; Tremblay and Schultz, 1999; Padoa-Schioppa and Assad, 2006; Kennerley et al., 2009; Rich and Wallis, 2014; Hunt et al., 2018; Jezzini et al., 2021; Stoll and Rudebeck, 2024b), and we hypothesized that the different attributes might specifically influence connectivity patterns. To investigate whether functional connectivity relies on the representations of specific decision variables, we examined connectivity between areas when we selectively removed decision-related representations from the firing rate of neurons in one of the two considered areas (see **Figure 3A** and **STAR Methods**). Here we focused on removing the influence of task-related activity in 12o neurons as area 12o showed high encoding of decision variables (**Figure 1**) as well as high level of connectivity with other ventral frontal areas (**Figure 2**). Note that we also applied similar methods on the activity of neurons from other areas as below. Briefly, we first conducted linear regression analyses containing multiple decision variables (chosen probability, flavor, and side for the stimulus period, reward and received flavor for the reward period) on the firing rate of each neuron and extracted the unexplained firing rate (residuals) from these models. We then correlated the residual activity of these 12o neurons with the unaltered activity of neurons from each other area using the previously described cross-validated CCA.

**Figure 3.**
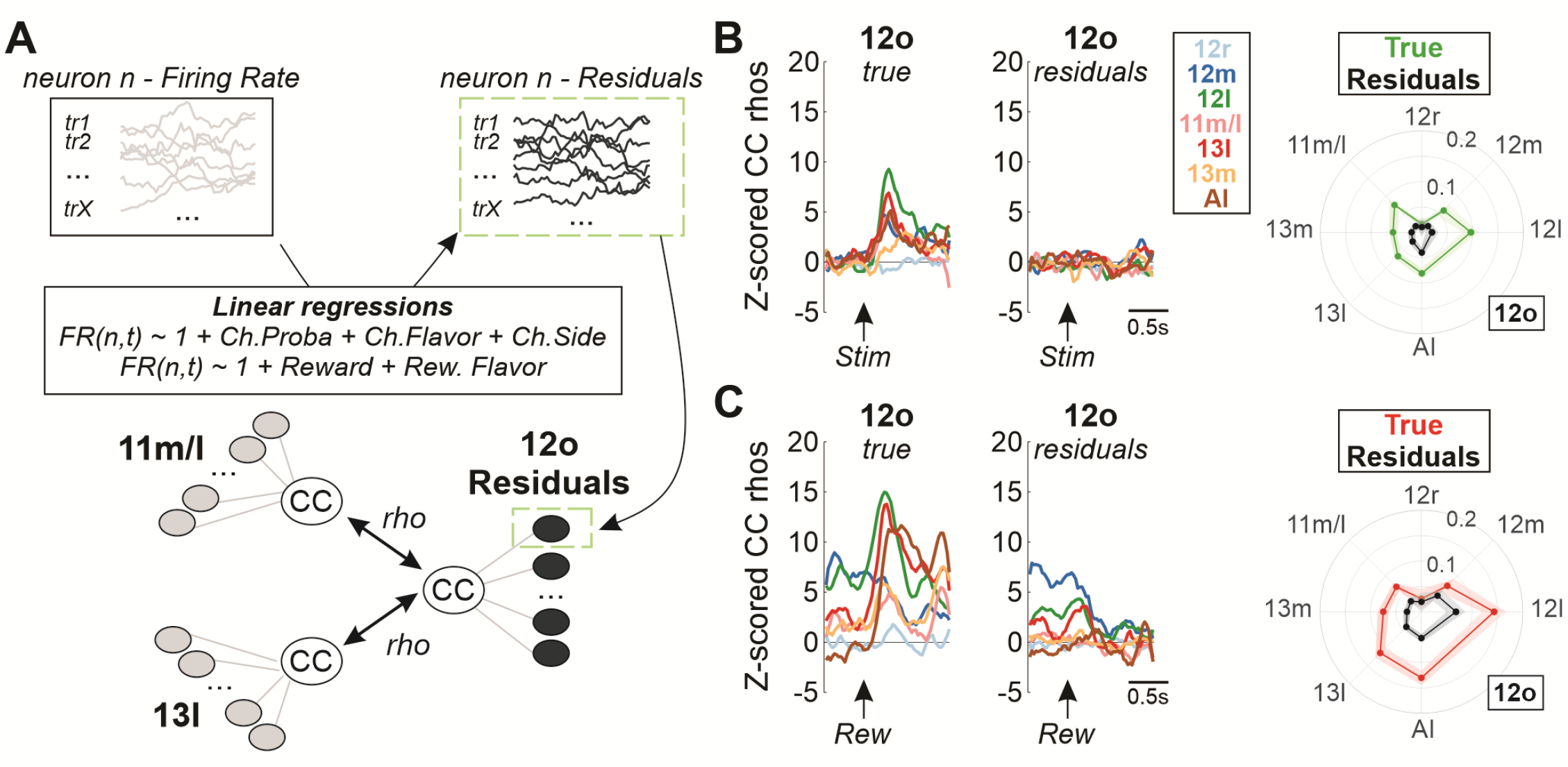
Altered connectivity following the removal of task-related activities. (**A**) Schematic of the procedure to assess the influence of specific representations on connectivity. Briefly, we first regressed out the activity related to behaviorally relevant decision variables (stimulus period: chosen probability, flavor and side; reward period: reward or no reward, reward flavor when delivered) from the firing rate of each 12o neuron across trials. We then applied CCA as previously described but now using the residual activity for 12o neurons (dark circles) and the unaltered activity for neurons of other areas (light circles). (**B**) Time course of the average zero-lagged correlations between 12o and other areas around stimulus onsets when using the true activity recorded in 12o (left, as in **Figure 2C**) or when correlating the residuals from 12o neurons obtained after regressing out the influence of decision variables (chosen probability, flavor and side) on each neuron’s firing rate. Right panel shows the average stimulus period connectivity fingerprint between the 2 conditions (green for true correlation, black when using residual activity in 12o). (**C**) Same as panel B but for reward onset. In that case, the residuals used for 12o were extracted from a two-way ANOVA model including whether a reward was delivered or not and the flavor of that reward when delivered (nested under reward). See also **Figure S2**.

We found that removing stimulus-related activities in 12o neurons largely abolished the increased connectivity we observed between 12o and all other areas following stimulus onset (**Figure 3B**). For instance, during the stimulus period the connectivity between 12o and all other areas remained at baseline levels (contrast left and right panels of **Figure 3B**). This decreased connectivity was in most part, but not exclusively, induced by the removal of chosen probability representations in area 12o compared to other decision variables (**Figure S2A**). A similar effect was observed following reward onset, where removing reward-related activity resulted in uniformly low connectivity during the reward period (**Figure 3C**). This was primarily driven by the encoding of the reward receipt compared to the flavor of the received reward (**Figure S2B**). Note that not all connectivity was abolished during the period preceding the reward. Such connectivity likely reflects a separate reward anticipatory process. Overall, this suggests that the connectivity between 12o and other ventral frontal areas predominantly, but not exclusively, relates to the prominent encoding of chosen probability and reward in this region (Stoll and Rudebeck, 2024b). Thus, the observed functional connectivity within ventral frontal cortex depended on specific task-related activity and likely reflects how information about the task is transferred from area to area during the decision-making process.

### Functional connectivity exists within specific activity subspace

A seminal study by Semedo and colleagues found that visual cortical areas in monkeys interact through specific communication subspaces of neural activity (Semedo et al., 2019). Importantly, this work revealed that two areas appeared to communicate through dedicated patterns of activity that were distinct from those used to communicate with other areas. Such ‘private’ communication channels between areas could represent a population-level mechanism through which specific information can be routed to different areas. This mechanism would reduce noise in interareal communication and allow interaction between areas to not only be specific but also to dynamically vary over time. Given the differences between frontal and sensory areas it remains an open question whether the functional interactions within ventral frontal cortex use specific communication subspaces between areas. Indeed, it is entirely possible that interareal communication between two or more other areas in frontal cortex use the same communication subspace as opposed to distinct subspaces for different interactions. The resulting communication between areas would be unspecific as all areas would share the same information to all other areas. Alternatively, and as proposed by Semedo and colleagues, an area could communicate with two or more areas through distinct communication subspaces.

To test these two possibilities in ventral frontal cortex, we conducted analyses to look for distinct communication between areas using two different approaches and looked for agreement between them to draw conclusions about whether areas were communicating through specific subspaces (**Figure 4A**). First, we compared the “direct” correlation strength between 2 areas as we described previously (e.g., correlation of 12o CC and AI CC obtained in the 12o-AI CCA, referred to as direct, *rdir*) with an estimated “indirect” correlation strength, *rind*, which represents the correlation between the canonical components extracted from 2 distinct CCAs (e.g., correlation of 12o CC obtained in the same 12o-AI CCA with the AI CC obtained from a different 13l-AI CCA). For the second approach, we correlated the CC of a given area obtained from CCA with 2 other areas (e.g., correlation of AI CC obtained in the AI-13l CCA with the AI CC obtained from the AI-12o CCA). We refer to this between-CCA correlation as *rbtw*.

**Figure 4.**
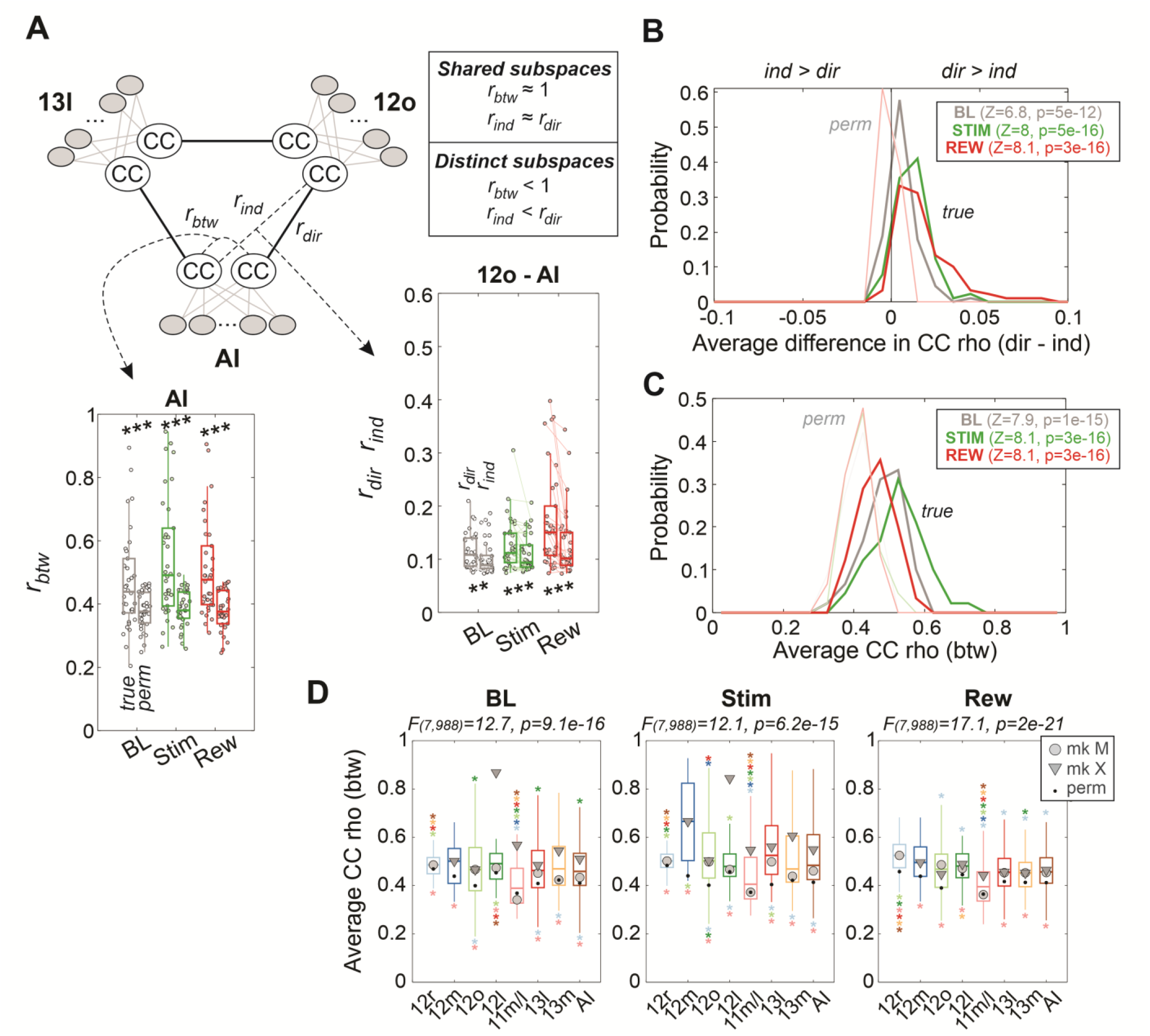
Functional connectivity between different pairs of areas exists within non-overlapping activity subspaces. (**A**) Schematic representation and example of the correlation measures extracted using cross-area CCA on neural activities from 3 areas, here 13l, 12o and AI. *r*_*dir*_ = absolute correlation strength between canonical components (CC) obtained from a unique CCA (similar to what was reported before). *r*_*ind*_ = absolute correlation strength between CC extracted across 2 distinct CCA (e.g., estimating the AI-12o correlation strength by correlating AI CC from AI-13l CCA with 12o CC from AI-12o CCA). If AI communicate with 13l and 12o using a shared activity subspace, then AI CC from AI-13l CCA should be similar to AI CC from AI-12o CCA, resulting in *r*_*dir*_ ≈ *r*_*ind*_. *r*_*btw*_ = absolute correlation strength between an area’s CC extracted by two distinct CCA (e.g., correlation between AI CC from AI-12o CCA and AI CC from AI-13l CCA). If AI communicate with 13l and 12o using a shared activity subspace, then both AI CCs should be perfectly correlated (*r*_*btw*_ ≈ 1). Bottom panels show example correlation strengths for all three measures. Stars represent statistical differences between *r*_*btw*_ and permutations (left panel) or between *r*_*dir*_ and *r*_*ind*_ (right panel) using one-tailed Wilcoxon signed rank test (** p<0.01 and *** p<0.001). (**B**) Distribution of the true average differences in CC correlations (*r*_*dir*_ - *r*_*ind*_) across all triplets of areas and for the three time periods (dark colors) compared to permutations (light colors, note that the permutations across time periods are overlapping). Reported statistics came from one-tailed Wilcoxon signed rank tests. (**C**) As panel B but showing the distributions of average between-CCA correlations (*r*_*btw*_). (**D**) Median (± 25 percentiles) between-CCA correlations (*r*_*btw*_) for each area across all possible triplets for both monkeys combined (boxplots) and individual monkeys (circles/triangles for monkey M/X respectively) and across the three periods of interest (baseline, stim and reward periods, from left to right). Black dots represent the median between-CCA correlations across permutations. Statistical significances (stars) were based on mixed-effect linear models assessing differences between areas. The location of the stars (p<0.05, FDR corrected) indicate the effect’s direction (e.g., red star – 13l – above the pink boxplot – 11m/l – means that 13l > 11m/l). BL: baseline; Stim: stimulus; Rew: reward periods

Our results across both methods support the idea that ventral frontal cortex subregions are interacting through distinct communication subspaces (**Figure 4**). For example, we found that AI communicated with its two neighboring areas 12o and 13l using different activity subspaces (**Figure 4A**). First, the direct correlation strength between AI and area 12o was significantly greater than when assessed using the canonical components from the AI-13l CCA (comparing *rdir* vs *rind*, **Figure 4A**, right). This means that the projected activity of AI neurons into the AI-13l communication subspace was not correlated with activity in 12o to the same degree as the best communication subspace between AI and 12o. Additionally, the between-CCA correlation strength (*rbtw*) of AI CCs from AI-13l and AI-12o communication subspaces was moderate, albeit significant, and overlapped with permutation strength in many sessions, suggesting a minimal overlap between communication subspaces between these areas. This was highly consistent when combining across all possible triplets of areas, where the direct correlation strength (indicative of a “private subspace” between areas) was always greater than the indirect method (**Figure 4B**) and where the between-CCA correlation was always significantly greater than permutations, although we note that the difference is slight (**Figure 4C**). Finally, some differences were observed in the between-CCA correlations across areas and periods of interest (**Figure 4D**). First, we observed variability between monkeys for areas 12l, 11m/l and 13m, with monkey X showing a higher level of subspace alignment than monkey M during the baseline and stimulus periods. Nevertheless, we found that area 11m/l exhibited the most distinct communication subspaces when communicating to other areas across all periods (**Figure 4D**; FDR-corrected post-hoc: 11m/l vs all other areas, W>3.15, p<4.7e-3 across periods).

Taken together, these analyses indicate that connectivity between two areas within ventral frontal cortex existed within specific activity subspaces, in that the canonical components in one area that best explained its relation to another area were largely distinct from those for other areas. Such a pattern indicates that an area communicates different information depending on the target area, a critical feature if each distinct subdivisions of ventral frontal cortex are representing and sharing information about distinct aspects of decision-making.

### Lag in functional connectivity between areas

The exchange of information between brain areas does not happen instantly but is influenced by factors such as synaptic efficiency and the speed of signal propagation along axons. Up to this point, we focused on zero-lag functional connectivity, in that CCA was applied on the simultaneously recorded activity of two areas. It is possible, however, that stronger functional connectivity could be observed when comparing activity recorded at different time points as there is a lag in the sending and receiving of information between areas. If connectivity from one area to another was higher at a different lag, this would be indicative of the directionality of information flow between areas. To explore this, we applied the same CCA method but in a cross-temporal manner, correlating the activity observed at one time point in one area with the activity observed at any other time point in another area. For each pair of areas, we quantified whether higher correlations were observed for negative or positive lags by computing a directionality index using the sum of the normalized correlation coefficients for lags up to 200ms (see **STAR Methods**).

Cross-temporal CCA revealed that the best correlation between areas was indeed not always observed at zero-lag (**Figure 5A**). For example, the connectivity between area 12l and 11m/l was not uniform across lags as there was a bias toward greater connectivity during the stimulus period when area 11m/l led area 12l (**Figure 5B**). Another notable lag existed between area 12o and AI, where the connectivity was greater when AI led 12o during the reward period. A similar trend was found between areas 13m and 12l, where area 13m led 12l during the reward period. All observed lags in connectivity are summarized on **Figure 6A**. It is important to note that the observed directionality is often unique to one of the time windows of interest. This suggests that the information flow across ventral frontal cortex is dynamic, and that a specific area might drive the activity of others during restricted time windows in relation with their specific role during reward-based decision-making. Indeed, removing the activity related to specific decision variables in neurons from area 11m/l not only reduced the overall connectivity with area 12l during the stimulus period but also partially removed the lag between these two areas (**Figure S3A**). A similar effect was observed when removing reward-related variables in AI, reducing the lag and overall connectivity with area 12o during the reward period (**Figure S3B**). Note that connectivity lags could nevertheless be observed when removing particular decision/reward-related activity, suggesting these lags could represent more general features of the network architecture.

**Figure 5.**
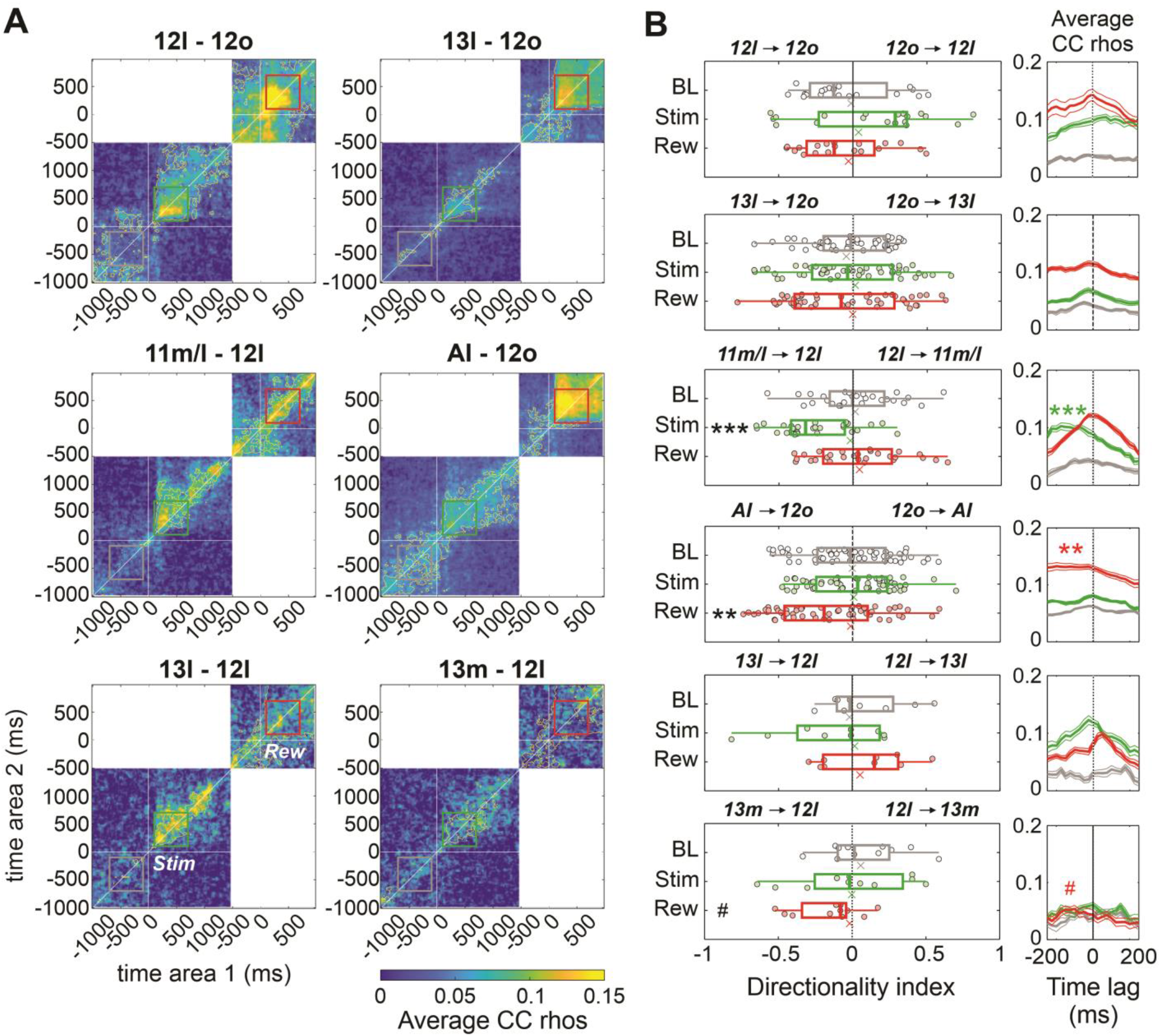
Cross-temporal CCA reveals the existence of lagged correlation between specific areas. (**A**) Examples of the averaged cross-temporal correlation matrices between canonical components extracted using cross-validated CCA for selected pairs of areas aligned on stimulus (bottom left matrices) and reward (top right matrices) onsets. Grey, green and red squares represent the time considered for the baseline (BL), stimulus (Stim) and reward (Rew) periods respectively. Yellow contours highlight time bins where a significant correlation was observed in more than 25% of the sessions (p<0.05, cluster corrected). (**B**) Left panels show the distribution of the directionality indices across sessions (dots) and periods of interest for the 6 example pairs of areas shown in panel A. A negative directionality index means that area 1 lead area 2, while a positive index means area 2 leads area 1. Stars indicate a statistical bias in the directionality index across sessions for a given period of interest (Wilcoxon signed rank test; # p<0.1, * p<0.05, ** p<0.01 and *** p<0.001). Right panels show the average canonical components correlation across the different time lags (between -200 and +200ms) for the 3 periods of interests, which form the basis of the directionality index.

**Figure 6.**
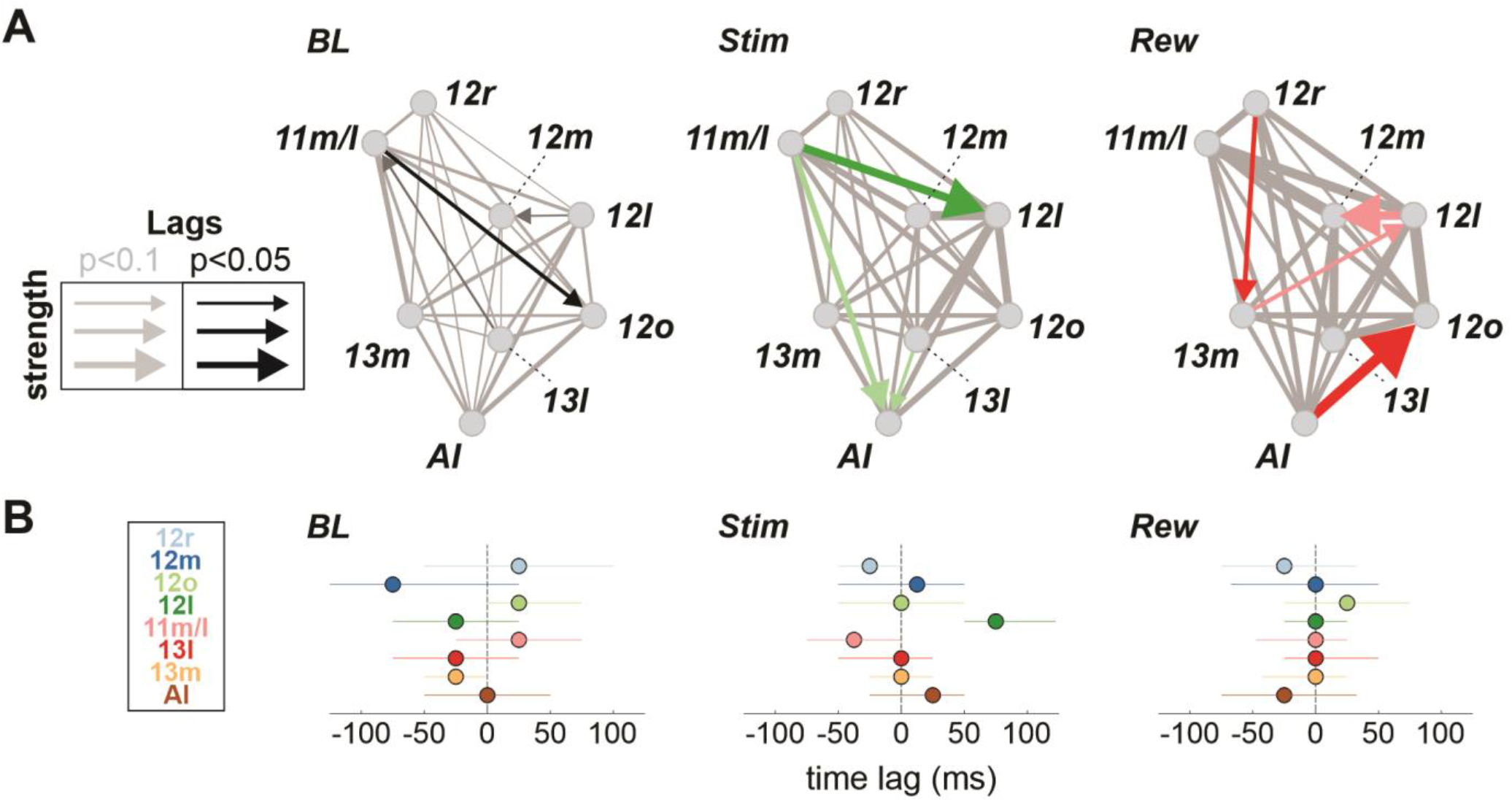
Lagged correlation across ventral frontal cortex. (**A**) Graph representation of the average correlation between canonical components across sessions during the baseline, stimulus onset and reward onset periods (left to right), at the time lag leading to the maximal correlation for each pair of areas. The thickness of each line represents the strength of the correlations (i.e., thicker lines for stronger correlations). Colored arrows represent a significant lag in the correlation between two areas (Wilcoxon signed rank test, light colors: p<0.1, dark colors: p<0.05). **(B)** Median latency [±10 percentiles] at which the maximal correlation was observed between a given area and all others computed for each session independently. A given pair of areas was not required to exhibit a significant lag to be included. See also **Figure S3**. BL: baseline; Stim: stimulus; Rew: reward periods.

Finally, we extracted the overall lag in communication by computing the median time lag between an area and all others across all neuronal populations considered (**Figure 6B**). No consistent lag was observed during the baseline period, which was characterized by high variability in the latencies of the peak correlation across sessions. During the stimulus period however, area 11m/l was more likely leading other areas while area 12l lagged behind (**Figure 6B**, middle panel). During the reward period, although the lags were more uniform across all areas, area 12o exhibited greater correlation after a lag. Notably, we previously found that area 12l exhibited the highest degree of integration of decision variable at the single neuron level during stimulus period while area 12o exhibited stronger representations during the reward period (Stoll and Rudebeck, 2024b). Altogether, our results support the idea that more integrative areas – area 12l during the stimulus period and area 12o during the reward – are receiving different streams of information from other ventral frontal area. Such a pattern of results would appear to indicate that these parts of area 12 have a more central role in distinct phases of the decision process.

## DISCUSSION

Using high-density single neuron recordings in monkeys performing a probabilistic choice task, we were able to reveal specific functional connectivity motifs across eight distinct cytoarchitectonic areas in ventral frontal cortex.

Specifically, areas 12o and 12l exhibited the strongest functional connectivity levels with other ventral frontal subdivisions during the stimulus and reward periods of our task, while area 13m was the least functionally connected (**Figure 2**). The observed connectivity was not only time-varying but 1) relied on the neural representations associated with the attributes of the different choice options that influenced subjects’ decisions (**Figure 3**) and 2) spanned unique population subspaces between pairs of areas indicating that sets of areas are communicating through specific channels (**Figure 4**). Finally, we identified markers of specific and directional information flow, notably with representations in area 11m/l leading those of areas 12l and AI during the stimulus period and representations in AI leading those of area 12o during the reward period (**Figures 5** and **6**). Taken together, our findings provide a novel insight into the dynamic functional communication across ventral frontal neuronal populations during decision-making, providing a functional account of how anatomically interconnected areas interact in higher cognitive function.

### 12o vs 12l are central nodes for reward-based decision-making

Our current work revealed that areas 12o and 12l are central nodes within ventral frontal cortex, exhibiting strong functional connectivity with many other subdivisions. Analyzing resting-state functional MRI data, Rapan and colleagues found only moderate levels of functional connectivity between area 12l and other ventral frontal areas (Rapan et al., 2023), consistent with what we observed during the baseline period. Following the onset of stimuli and reward delivery, 12l connectivity greatly increased, especially with areas 12o, 11m/l and 13l. The difference in connectivity between baseline/resting-state and behaviorally engaged periods suggests that functional connectivity depends on the processes that are currently engaged, in this case reward-based decision-making. Anatomically, both area 12l and 12o areas are strongly interconnected with most other subdivisions of ventral frontal cortex (Carmichael and Price, 1996). However, their connectivity diverges in that area 12l exhibits dense connectivity with lateral frontal areas 45 and 46v, while area 12o exhibits greater connectivity with medial frontal areas (Carmichael and Price, 1996; Saleem et al., 2014; Rapan et al., 2023). This dissociation is likely critical to the function of both areas. In fact, our previous work highlighted major differences in how different ventral frontal subdivisions represented decision variables, with area 12l exhibiting diverse and integrative encoding of decision variables during the stimulus period while area 12o showed a more unique pattern of activity related to outcome probability and rewards (Stoll and Rudebeck, 2024b). Combined with our finding that representations in areas 12l and 12o lagged behind other subdivisions during the stimulus and reward periods, respectively, our results indicated that areas 12l and 12o may play distinct integrative roles in decision-making and potentially other cognitive processes. Furthermore, we found that removing the influence of decision/reward-related variables greatly reduced the connectivity across the whole network. Indeed, specifically removing the encoding of probability from the activity of 12o neurons was the most effective way to reduce 12o connectivity with other parts of ventral frontal cortex compared to removing the influence of flavor or side information. This strongly supports the notion that the observed functional connectivity between areas relates to the representation of specific decision variables, in this case probability. It also highlights that resting state functional connectivity does not necessarily reflect the diversity of connectivity that networks might be able to display.

### Stimulus-related connectivity

During the stimulus period, we found that the neural representations in area 11m/l led those in AI and area 12l. Such a temporally specific connectivity pattern fits with the proposed role of area 11m/l in guiding goal selection through the representation of stored specific stimulus-reward value associations (Murray et al., 2015). Related to this, our prior work found strong representations of stimulus-flavor associations in area 11m/l and 12l compared to more posterior parts of ventral frontal cortex just after the visual stimuli were presented in the trial (Stoll and Rudebeck, 2024b). It is possible that the lag in connectivity observed here could represent the broadcast of this stimulus-flavor information to other areas like area 12l where it is integrated with other attributes of the available options to adaptively guide the choice. Counter to this idea, we found that removing the variance in neural activity related to juice flavor in area 11m/l did not eradicate this lead in connectivity between this area and 12l, nor did it remove all connectivity with area 12l. It is possible that our analytic approach to isolate decision-related information does not fully remove all influence on neural activity (notably non-linear or time-varying representations) or that other variables not captured by our models are transmitted through this connectivity. Alternatively, such pattern might suggest that the network is biased toward representations in area 11m/l leading those in more posterior/lateral areas, irrespective of which information is transmitted. Evaluating the connectivity between area 11m/l and 12l in during other cognitive processes will help to understand the diversity of representations that can be shared through this channel.

### Reward-related connectivity

We found that AI representations led those in area 12o during the period where the reward was delivered. Prior tract tracing work has shown that AI sends dense connections to all posterior ventral frontal cortex subdivisions including areas 13m, 13l, and 12o, but sends few to area 12l (Carmichael and Price, 1996). AI receives primary sensory information including visceral and gustatory (Ongur and Price, 2000) and is contiguous with the more posterior insula cortex which has been hypothesized to represent internal states (Craig, 2002, 2009; Evrard, 2019). The strong lagged connectivity we observed between AI and area 12o could underlie such function, enabling the updating of stimulus-reward probability associations depending on whether reward was delivered. This would fit previous reports linking the activity in area 12o with information seeking and contingent learning by assigning rewards to chosen stimuli (Chau et al., 2015; Folloni et al., 2021; Jezzini et al., 2021).

Less clear is why we failed to find strong connectivity between AI and area 13m given the roles of these two areas in updating sensory-specific valuations and internal states (Rudebeck and Murray, 2014; Padoa-Schioppa and Conen, 2017). One potential reason could relate to the precise location of our AI recordings. AI is composed of distinct subdivisions, each of which exhibit specific patterns of anatomical projections to other areas ventral frontal areas (Carmichael and Price, 1996). In particular, the intermediate AI (Iai) sends heavy projection to area 12o but not 13m/l, while lateral and posterior AI (Ial, Iapm and Iapl) show the opposite pattern. Although we did not assess the precise boundaries between subdivisions of AI in our current work, our recordings were more heavily concentrated in the most anterior portions of AI, specifically intermediate AI. This means that we were more likely to capture interactions between AI and 12o than with other subdivisions. Aside from this, it is also possible that our task was not diverse enough to truly capture the information flow from/to area 13m, which might be more related to learning and updating the sensory-specific values associated with a stimulus (Murray et al., 2015). On this view, as the juice flavors were stable and did not change in each session, there was no need to update juice reward associations and so connectivity between the areas was low. Interestingly, we did observe an increase in connectivity between area 13m and neighboring subdivisions later in each trial, around the inter-trial interval. This connectivity was temporally dissociated from outcome flavor representations observed in this area, but matched the late emergence of reward responses in area 13m compared to other subdivisions (Stoll and Rudebeck, 2024b). Such temporal patterns fit a role in value updating processes, wherein that connectivity could be a marker of updating internal states that might occur later after the reward on each trial has been delivered. Taken together, functional connectivity between area 13m and other ventral frontal subdivisions is likely to be engaged when learning is experimentally modulated or when internal states change over the course of a session when animals become sated.

## Limitations

Inter-areal functional connectivity within ventral frontal cortex has been assessed using functional MRI, notably while subjects are not behaviorally engaged in a task (e.g., (Kahnt et al., 2012; Rapan et al., 2023)). Although such an approach continues to provide insights into brain-wide interactions, signal distortion within ventral frontal cortex likely influences the overall detection of functional connectivity (Devlin et al., 2000) and the extent to which the observed connectivity relates to underlying neural activity remains unclear (Leopold and Maier, 2012). Here, we took the complementary approach of assessing local functional connectivity using the activity of single neurons recorded simultaneously across many subdivisions of ventral frontal cortex. Such an endeavor is limited by the ability to record from many areas simultaneously and consequently our study used relatively small populations of neurons. Nevertheless, connectivity estimates were reproducible across the numerous sessions and distinct populations of neurons, giving credence to our observations. Recent advances in recording techniques will soon make it possible to record neural activity from many more areas simultaneously (Steinmetz et al., 2021; Jung et al., 2024; Liu et al., 2024), a critical feature to further our understanding of the complex network dynamics underlying decision-making processes.

Also, we found that most interactions within ventral frontal cortex peaked at zero-lag, occurring almost instantaneously. Surprising at first, such zero-lag correlations are likely a byproduct of the window sizes and Gaussian smoothing we applied to neurons’ activity. Neurons in frontal areas often exhibit low firing rates, necessitating the use of sliding window analyses with large bin sizes and/or Gaussian smoothing kernels (Kass et al., 2003). The drawback of such approaches is that they limit temporal resolution. Direct synaptic delays between neighboring areas in ventral frontal cortex are likely shorter than the 25ms bin size used here and thus areas appear to be interacting instantaneously. Such feature also relates to the well-known limitation of connectivity analyses, including CCA, where the observed correlations might not exclusively reflect the direct communication between two areas. Such correlations could be mediated through a third area or reflect the influence of a common input. Causally manipulating specific pathways while assessing connectivity would provide insight into whether measures of functional communication between two areas are direct or not.

Finally, the exact nature of the information being transmitted between two areas remains unclear. Although we found that removing firing rate modulations related to specific decision variables greatly reduced the observed functional connectivity, further modeling is required to pinpoint the precise information being exchanged. For example, one could reproduce the activity of our neuronal populations using multi-area recurrent neural networks and extract the dynamical interactions across areas when artificially lesioning specific part of the network (e.g., (Perich et al., 2020)).

## Conclusion

Careful anatomical work revealed that subdivisions of ventral frontal cortex form a heavily interconnected network (Carmichael and Price, 1996; Rapan et al., 2023). Functionally, neurons within distinct ventral frontal subdivisions exhibit highly diverse and selective representations, subserving distinct functions during reward-based decision making (Stoll and Rudebeck, 2024b). Interconnected areas, however, don’t function in isolation. Our current study reveals that interactions between neuronal populations across ventral frontal cortex are spatially and temporally specific during decision-making. It further emphasizes the possible existence of what appear to be subdivisions, namely 12o and 12l, where signals are directed to be integrated into distinct processes. Our study therefore provides a unique functional account of the previously described anatomical networks spanning these areas. Characterizing how this network of areas interacts during a diverse set of cognitive and affective functions, and how these areas interact within the larger frontal networks that include medial and lateral frontal cortex (Carmichael and Price, 1996) will be critical to our understanding of the neural dynamics supporting behavior.

## ACKNOWLEDGEMENTS

This work was supported by a National Institute of Mental Health BRAINS award to PHR (R01s MH110822; MH132064), a young investigator grant from the Brain and Behavior Foundation (NARSAD) to PHR, a Philippe Foundation award to FMS, seed funds from the Icahn School of Medicine at Mount Sinai to PHR. We thank Marques Love and Dr. Patrick Hof for their help in defining neuroanatomical boundaries.

## AUTHORS CONTRIBUTIONS

Conceptualization and Methodology: FMS, PHR; Investigation, Data curation, Analysis, Software and Visualization: FMS; Writing – Original Draft and Review/Editing: FMS, PHR; Funding Acquisition and Supervision: PHR.

## STAR METHODS

### RESOURCE AVAILABILITY

#### Lead contact

Further information and requests for resources should be directed to and will be fulfilled by the lead contact, Frederic M. Stoll.

#### Materials availability

This study did not generate new unique reagents.

#### Data and code availability

- Behavioral and neurophysiological data reported in this paper will be shared by the lead contact upon request.
- All original code will be deposited and publicly available at https://github.com/RudebeckLab/POTT-conn before publication.
- Any additional information required to reanalyze the data reported in this paper is available from the lead contact upon request.

### EXPERIMENTAL MODEL AND SUBJECT DETAILS

Subjects were two adult male rhesus macaques (*Macaca mulatta*), monkeys M and X, aged 8 and 5.5 years old, and weighing 11.9 and 7.9 kg, respectively, at the start of the neurophysiological recordings. Animals were grouped-housed, kept on a 12-h light/dark cycle and had access to food 24 hours a day. Throughout training and testing each monkey’s access to water was controlled for 5 days per week. All procedures were reviewed and approved by the Icahn School of Medicine Institutional Animal Care and Use Committee.

## METHOD DETAILS

The dataset analyzed here has been discussed in previously published work. Notably, it was used to assess how single neuron responses integrated taste preference differentially across prefrontal-limbic circuit (Stoll and Rudebeck, 2024a) and how neurons within specific anatomically-defined subdivisions of OFC and vlPFC encoded decision variables in a dissociable and dynamic manner (Stoll and Rudebeck, 2024b). No attempts were made at the time to understand how neurons’ activities relate to one another, which is the focus of the current set of analyses. We describe the novel analyses in the following sections, although we refer readers to these previous works for additional methodological details which we may only briefly describe here.

### Apparatus

Monkeys sat with their heads restrained in a custom primate chair situated 56 cm from a 19-inch monitor screen. Gaze location was monitored and acquired at 90 frames per second using an infrared oculometer (PC-60, Arrington Research, Scottsdale, AZ) and used to report choices. Juice rewards were delivered to the monkey’s mouth using custom-made air-pressured juice dispenser systems (Mitz, 2005). Trial events, reward delivery, and timings were controlled by MonkeyLogic (NIMH, version 1) behavioral control system, running in MATLAB (version 2014b, The MathWorks Inc.). Raw electrophysiological activity was recorded at 40kHz resolution using an Omniplex data acquisition system (Plexon, Dallas, TX). Spikes from putative single neurons were automatically clustered offline using the MountainSort plugin of MountainLab (Chung et al., 2017) and later curated manually based on principal component analysis, inter-spike interval distributions, visually differentiated waveforms, and objective cluster measures (Isolation > 0.75, Noise overlap < 0.2, Peak signal to noise ratio > 0.5, Firing Rate > 0.05 Hz). Details on the isolation quality of the single neurons can be found in the previously published work on this dataset (Stoll and Rudebeck, 2024a).

### Behavioral task

During recording sessions, monkeys performed 3 closely related tasks: Single option, Instrumental and Dynamic probabilistic tasks (Stoll and Rudebeck, 2024a). Our analyses here focused on instrumental trials (**Figure 1**). In this task, monkeys could choose between two options presented simultaneously on the right and left side of the screen. Each option was composed of two features: an external-colored rectangle indicating which outcome flavor (out of 2 possible juices per session) monkeys could earn and a central rectangle, more or less filled, indicating the probability at which this particular outcome flavor would be delivered at the end of the trial. Monkeys were faced with options containing one of two possible colors on a given session (randomly selected from a set of 9 colors) associated with two different juice flavors (randomly picked from a set of 5, which included apple, cranberry, grape, pineapple and orange juices, diluted in 50% water). Probabilities used were from 10% to 90% (by steps of 20% for monkey M and 10% for monkey X).

Trials were initiated by fixating a central fixation cross for 0.7 to 1.3s (steps of 0.3s), at which time monkeys were free to look at the two options (displayed for 0.4 to 0.8s by steps of 0.2s, pseudorandomly selected). Stimuli were then turned off for 0.2s and two response boxes appeared on both sides of the previously shown options (3 possible locations equidistant to the options’ locations; bottom left/right, center left/right, top left/right). Monkeys had to fixate the response box on the side of the desired option within 8s to make their choice. They had to maintain fixation on the selected response box for a minimum of 0.25s to register the response, at which time the other response box would disappear. The requirement for them to maintain their gaze on one of the options meant that subjects could modify their choice within a trial if they so choose. Continued fixation was required for an additional 0.6 to 1.2s (steps of 0.3s), after which the response boxes would disappear for 0.3 to 0.7s (steps of 0.2s). Both options were presented again at the same locations (feedback period), with the selected one initially flashing (5 times 0.1s ON followed by 0.1s OFF, total time of 0.5s), before staying on the screen for the duration of the reward (if delivered) and an additional 0.5s. In rewarded trials, monkeys received 2-3 pulses of 0.03-0.06s of fluid (separated by 0.1s each, 0.25-0.36ml total reward per trial of the outcome flavor and at the probability indicated by the selected option). Nonrewarded trials were matched in time to trials that included reward delivery. Finally, rewarded trials were followed by a 2s intertrial interval, and unrewarded trials were followed by 3.5-4s. If monkeys failed to maintain fixation when required, a large red circle was presented at the center of the screen for 1s, followed by a longer intertrial interval (4-6s for monkey M, 3-4s for monkey X). This was done to ensure that subjects received feedback about their erroneous actions. Failure to initiate a trial by looking at the fixation star within 6s of its appearance resulted in the same red circle and intertrial interval.

The two options were associated with different outcome flavors (juice 1 vs. 2) in half of the trials for monkey M and in 3 out of 4 trials for monkey X. In these trials, the probability was always different for the two outcomes in monkey M but could be either different or similar in monkey X (e.g., juice 1 at 70% vs. juice 2 at 70%). In the remaining trials (1/2 for monkey M and 1/4 for monkey X), the two options were associated with the same outcome flavor (e.g., juice 1 vs. juice 1) but with different probabilities. Both trial types were considered in the following analyses except when assessing the role of decision variable encoding on functional connectivity (see *“CCA on residual activity”*).

### Surgical procedures and neural recordings

During aseptic surgeries, in a dedicated operating theater, under full anesthesia, and while being constantly monitored, monkeys were implanted with a titanium head restraint device and a form-fitted PEEK recording chamber that contained a 157-channel semi-chronic microdrive system (Gray Matter Research, Bozeman, MT; **Figure 1**) housing glass-coated electrodes (1-2MΩ at 1kHz; Alpha Omega Engineering, Nazareth, Israel). Cranial implants were held in place using orthopedic grade titanium screws and small amounts of dental acrylic (C&B Metabond, Parkell Inc, Edgewood, NY; Ortho-JET BCA, Lang Dental MFG Co., Wheeling, IL). Recording locations were confirmed using several approaches. First, we recorded the cumulative depth of each electrode when slowly lowering them, tracking changes in background noise and electrophysiological activity suggesting white/gray matter transitions. We also acquired CT images at different time points, which were co-registered to post-operative MRIs. Finally, we captured and digitalized block-face pictures before cutting every brain section which were later stained, showing clear marks of the electrodes’ track. Combined with microscope observations of histological stained sections (see below for more details) and using Free-D software (Andrey and Maurin, 2005), this allowed us to reconstruct the precise anatomical location of every electrode and recorded neurons.

### Tissue preparation and immunohistochemistry

Following recordings, monkeys were deeply anesthetized and transcardially perfused with 4% formaldehyde in Phosphate-buffered saline. The brain was extracted, post-fixed and cryo-preserved before being shipped to FD NeuroTechnologies, Inc (Columbia, MD) for tissue preparation and staining. Briefly, the recorded brain hemispheres were further cryoprotected in solution before fast freezing in isopentane. Serial sections of 50μm were cut coronally and series of 4 consecutive sections were collected and stained separately (resolution of 200μm per staining). Every first section of each series was stained using cresyl violet solution (Nissl staining), while the second and third series were processed for Calbindin (using mouse monoclonal anti-Calbindin-D-28K antibodies, 1:1000 dilution; Millipore Sigma, St. Louis, MO) and SMI-32 (using mouse Purified anti-Neurofilament H nonphosphorylated antibodies, 1:12000 dilution; Biolegend, San Diego, CA) immunohistochemistry.

### Defining neuroanatomical boundaries

We looked for variation in the composition of cortex across the ventral frontal cortex of each of our subjects on the Nissl, SMI-32, and calbindin-stained sections, following the approach from Carmichael and Price (Carmichael and Price, 1994) and later supported by Rapan and colleagues (Rapan et al., 2023). Sections were inspected using either Nikon or Zeiss light microscopes. In each monkey we were reliably able to discern areas 13l, 13m, 12m, 12r, 12l, and 12o. While areas 11m and 11l could be differentiated in one subject, they were harder to identify in the other. We therefore combined these two areas into one which we label 11m/l. We did not attempt to discern subdivisions of AI as the number of neuronal populations in each would have been too few to analyze, although we note that our recordings mainly targeted the most anterior sections of AI. We refer the readers to our previous work for the precise definitions of each area and examples of areal boundaries from this dataset (Stoll and Rudebeck, 2024b).

### QUANTIFICATION AND STATISTICAL ANALYSIS

#### Pre-processing of neurophysiological data

Neurons were included in the analysis based solely on the quality of isolation and firing rate. Their functional response patterns were not taken into account. Spiking activity for each trial was first smoothed using a 25ms Gaussian kernel before being averaged over 25ms bins. Preliminary analyses using longer bin sizes led to similar observations and are not reported (see also Discussion). Neurons’ firing rates were aligned to multiple events across trials (central fixation, stimulus onset, response fixation, feedback and reward onsets). Our analyses were performed at a 25ms time resolution before being averaged around events of interest, namely the baseline (100-700ms before stimulus onset), stimulus (100-700ms following stimulus onset) and reward (100-700ms following reward delivery) periods. We rejected neurons with an average firing rate across trials and time bins (around stimulus and reward onsets) lower than 0.5Hz. This threshold and the initial smoothing of firing rates was required for the following analyses to converge. Results are provided for each monkey as well as combined.

### Functional interaction using Canonical Component Analysis (CCA)

We investigated the population interaction between pairs of areas using canonical correlation analysis (CCA) (Härdle and Simar, 2007; Semedo et al., 2020). Briefly, CCA isolates a set of dimensions, referred to as canonical components, contained within the population activity of two areas such that these dimensions maximally correlate. In our study, we only considered the first canonical component. We analyzed sessions with at least 4 neurons simultaneously recorded in each of the two areas considered. All pairs of areas were considered but only pairs of areas with at least 3 replicates (a replicate being a single session in which we recorded enough neurons in the 2 considered areas) were further analyzed (**Table S2**). Indeed, we could not assess the connectivity between 12r and 12m as we did not consistently record enough neurons simultaneously in these areas. To avoid overfitting issues, we performed CCA using a 10-fold cross-validation procedure. Specifically, we first extracted the weight associated with each neuron across the trials used in the training set (*canoncorr* function in Matlab) and then used them to correlate the activity of neurons in the testing set, following:

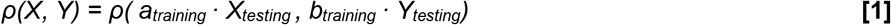

where *ρ* is the canonical correlation coefficient, *X* and *Y* the matrices containing the responses of simultaneously recorded neurons across trials from the testing sets and at a given time, and *a* and *b* the weights for neurons from area X and Y, respectively, extracted using the training set. We reported the average correlation coefficient across the 10-folds.

To assess the temporal evolution of population activity correlations, we applied CCA at equivalent time points (by extracting the canonical components of two areas at a given time, referred to as zero-lagged correlations) but also at different times (by correlating the population activity of one area at time *t* with the activity of another area at time *t±i*, referred to as cross-temporal CCA, see **Fig. 5**). Such procedure has the potential to highlight whether specific population activity might lead other patterns of activity. All lags were tested for both event alignments (stimulus onset: -1000 to 1500ms by steps of 25ms, 100 bins; reward onsets: -500 to 1000ms by steps of 25ms, 60 bins). We computed the significance of the observed correlation coefficients by permuting once the trial order of one of the considered neuronal populations at every time bin and at every lag. This resulted in 13,600 CCAs across time per pair of areas. Permuted correlation coefficients were uniform across time, which allowed us to use all permutations across time bins to assess the significance of true cross-temporal correlations. We used cluster-based corrections with a cluster-defining threshold at p<0.05 and a cluster size of at least 10 time-bins (*bwconncomp* function in Matlab, ignoring corners by using connectivity = 4).

Within periods of interest (100 to 700ms following stimulus and reward onsets), we computed the average correlation coefficients for lags up to ±200ms, on which we could extract a directionality index as follows:

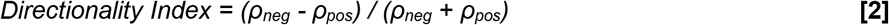

where *ρneg* and *ρpos* represent the sum of the min-max normalized correlation coefficients for negative and positive lags, respectively. We assessed significance in directionality index by comparing it across sessions to the matching permuted index for each considered pair of area from our permutation using Wilcoxon signed-rank tests.

### CCA on residual activity

To investigate whether the observed connectivity was related to the content of the information encoded, we computed the zero-lagged correlation under normal condition (as described before) as well as after removing the influence of the various stimulus- and reward-related parameters. This was performed by first explaining the activity of every neuron from one area using ANOVAs. We limited this analysis to the response of neurons during trials where different outcome flavors were offered, as previously reported (Stoll and Rudebeck, 2024a, 2024b). Stimulus-related activity was first explained using a three-way ANOVA which included the chosen probability (10-30-50-70-90%, linear), chosen flavor (J1 or J2, categorical) and chosen side (left or right, categorical) as factors. We also used one-way ANOVAs which included each of the three decision-related factors to explain stimulus-related activity. Reward-related activity was explained using two-way nested ANOVAs which included reward receipt (delivered or not, categorical) and the received flavor when rewarded (J1 or J2, categorical) as factors. As before, we also used one-way ANOVAs with reward receipt or flavor received (in rewarded trials only) to explain reward-related activity. In all cases, we then extracted the residual activity (i.e., the unexplained variance in firing rate) at every time bin. We used the residual activity for each neuron from a given area and extracted the maximal correlation with the full activity of neurons from another area using CCA, as previously described. We compared the z-scored correlations depending on whether we used the raw activity or the different residual models using mixed-effect linear regression, which included the condition (5 categorical levels for stimulus period: true correlation, three-way ANOVA model, chosen probability model, chosen flavor model and chosen side model; 4 categorical levels for reward period: true correlation, two-way nested ANOVA model, reward model, and rewarded flavor model), and areas (7 levels) as main factors, as well as monkeys and sessions as random effects. Here, we mainly focused on the correlation between the activity of 12o neurons and neurons from other areas, as area 12o showed strong connectivity during stimulus and reward periods. This meant that we applied ANOVAs and extracted residual activities only for 12o neurons (see **Figure S3** for this method being used on different pairs of areas). Thus, this approach enables us to isolate whether the observed correlations were related to the representation of decision-related variables.

### Cross-area CCA

The above analyses enabled us to determine if there is a correlation in activity between two areas. The parts of the ventral frontal cortex that we have obtained recordings from are, however, highly interconnected (Carmichael and Price, 1996; Rapan et al., 2023). Given this we wanted to understand whether a given area might communicate with two other areas using distinct neural subspaces. If this was the case it would potentially indicate that specific information was being transferred between a specific pair of areas. To assess whether the observed connectivity relied on similar or distinct subspaces, we considered triplets of areas recorded simultaneously, while keeping other requirements similar (e.g., minimum number of neurons or replicates). In brief, we extracted the canonical components (CC) from CCAs applied to all possible pairs of areas within each triplet (i.e., area X – Y, area Y – Z and area X – Z). We then used the weights for a given area obtained from one of the CCA and assessed the correlation coefficients when the other area was assigned the weight coming from a different CCA (see example in **Fig. 4A**). Specifically, we extracted the absolute maximal canonical correlations for each set of three areas, which we referred as *ρdir*, as previously described:

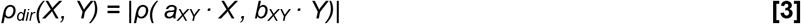

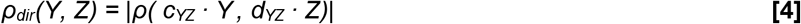

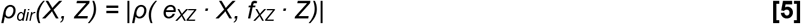

where *a, b, c, d, e* and *f* are the weights for neurons recorded in areas *X, Y* and *Z*. We then derived the indirect canonical correlation (*ρind*) between 2 areas, X and Y for example, using the canonical components obtained from two independent CCAs (X vs Y and Y vs Z), using:

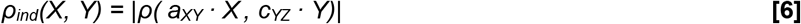

We then compared *ρdir* and *ρind* for each pair of areas. Lower correlation coefficients for *ρind* compared to *ρdir* indicated that area Y communicated with areas X and Z through different neural subspaces. We also assessed the correlation between the canonical components within a single area, using:

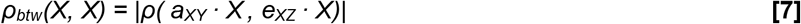

In this case, a low *ρbtw* correlation coefficient would indicate that area X communicated with areas Y and Z using different neural subspaces.

This procedure was 10-fold cross-validated and computed for zero-time lag only. It is important to note that we used here the absolute value of the correlation coefficients before averaging across the 10-fold cross-validation procedure. Such a transformation was necessary as the sign of the neurons’ weights across two independent CCAs are arbitrary, which could result in negative correlations if they were opposite by chance. To alleviate the potential bias toward higher correlation values, we compared the true average correlations with 100 permutations, following the same procedure. In this case, we permuted the trial order of two out of the three neural populations. Average permuted correlations were indeed greater than 0 due to taking this transformation.

## SUPPLEMENTARY INFORMATION

**Figure S1.**
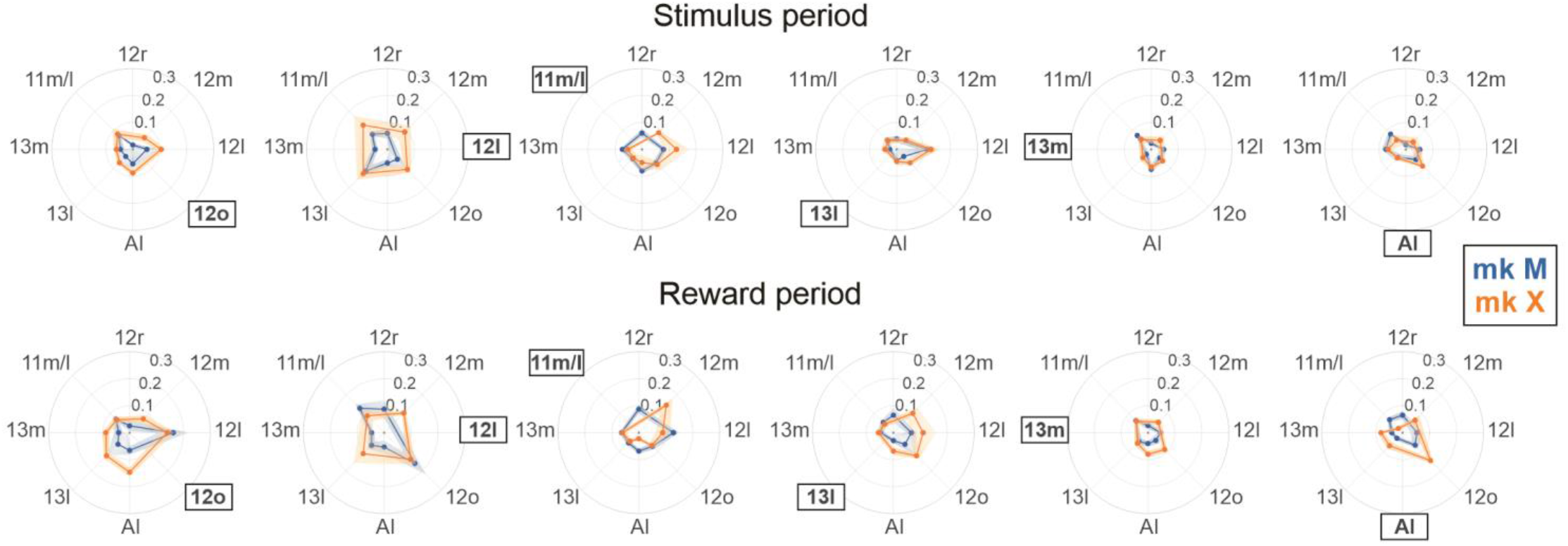
Connectivity fingerprints across monkeys. Related to Figure 2. Connectivity fingerprints across areas for population of neurons recorded in monkey M (blue) and X (orange) during the stimulus (top panels) and reward (bottom panels) periods. Fingerprint maps for area 12r and 12m are not shown given the low number of sessions when considering individual monkeys.

**Figure S2.**
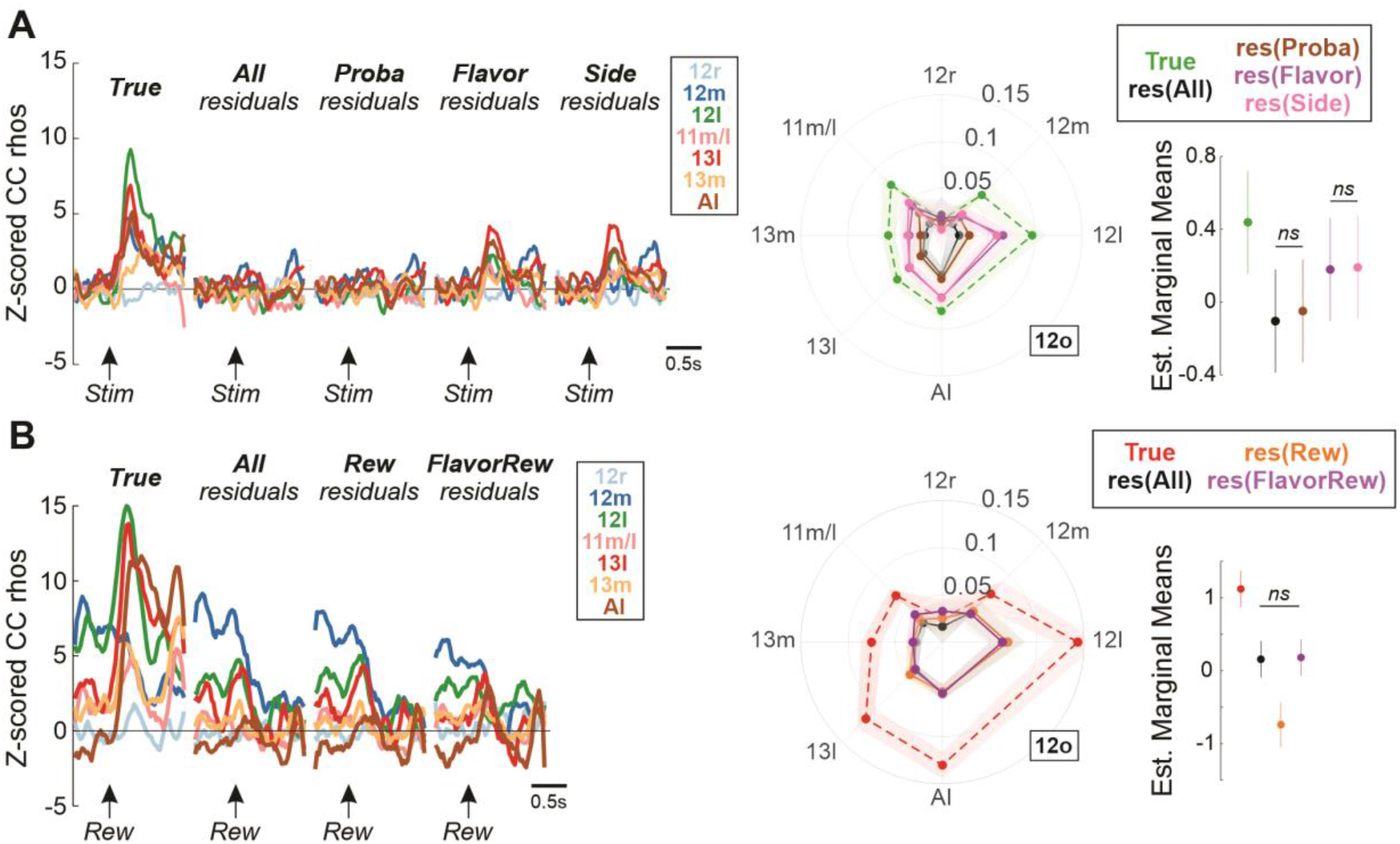
Altered connectivity following the removal of specific task-related activities. Related to Figure 3. (**A**)Same as **Figure 3** but the residuals used for 12o were also extracted from one-way ANOVA models including a single variable of interest (chosen probability or chosen flavor or chosen side). “True” and “all residuals” conditions are reproduced from **Figure 3B**. Middle panel shows the average stimulus period connectivity fingerprint across the 5 different conditions (green for true correlation, black when using the residual activity of a model including the 3 variables of interest, brown when the model included only chosen probability, violet when the model included only chosen flavor, and pink when the model included only chosen side). Right panel show the estimated marginal means extracted from a mixed-effect linear model explaining the Z-scored correlations using conditions (5 levels) and areas (7 levels) as main factors, and monkey/sessions as random effects. We found significant decreases in connectivity between 12o and all other areas when using the residuals from either models compared to the true connectivity (mixed-effect linear regression, factor lesion type, F_(4,1037)_=23.3, p=1.6e-18; FDR-corrected post-hoc: true vs all/proba/flavor/side residuals, W>3.9, p<1.7e-4). A stronger decrease following the removal of chosen probability was observed (FDR-corrected post-hoc: proba vs flavor/side residuals, W>3.5, p<4.2e-4; flavor vs side residuals, W=0.18, p=0.85), to a similar degree than what was observed when removing all considered variables (FDR-corrected post-hoc: proba vs all residuals, W=0.87, p=0.42). (**B**)Same as panel A but for the reward period, where models included reward (delivered or not, orange), the delivered reward flavor (*FlavorRew*, violet, only in rewarded trials) or both (*All*, with reward flavor nested under reward, black). As before, we found a significant decreases in connectivity when using the residuals from either models compared to the true connectivity (mixed-effect linear regression, factor lesion type, F_(3,831)_=51.5, p=1.4e-30; FDR-corrected post-hoc: FlavorRew vs all, W=0.25, p=0.79, other comparisons, W>6.8, p<2.1e-11). Rew: reward period.

**Figure S3.**
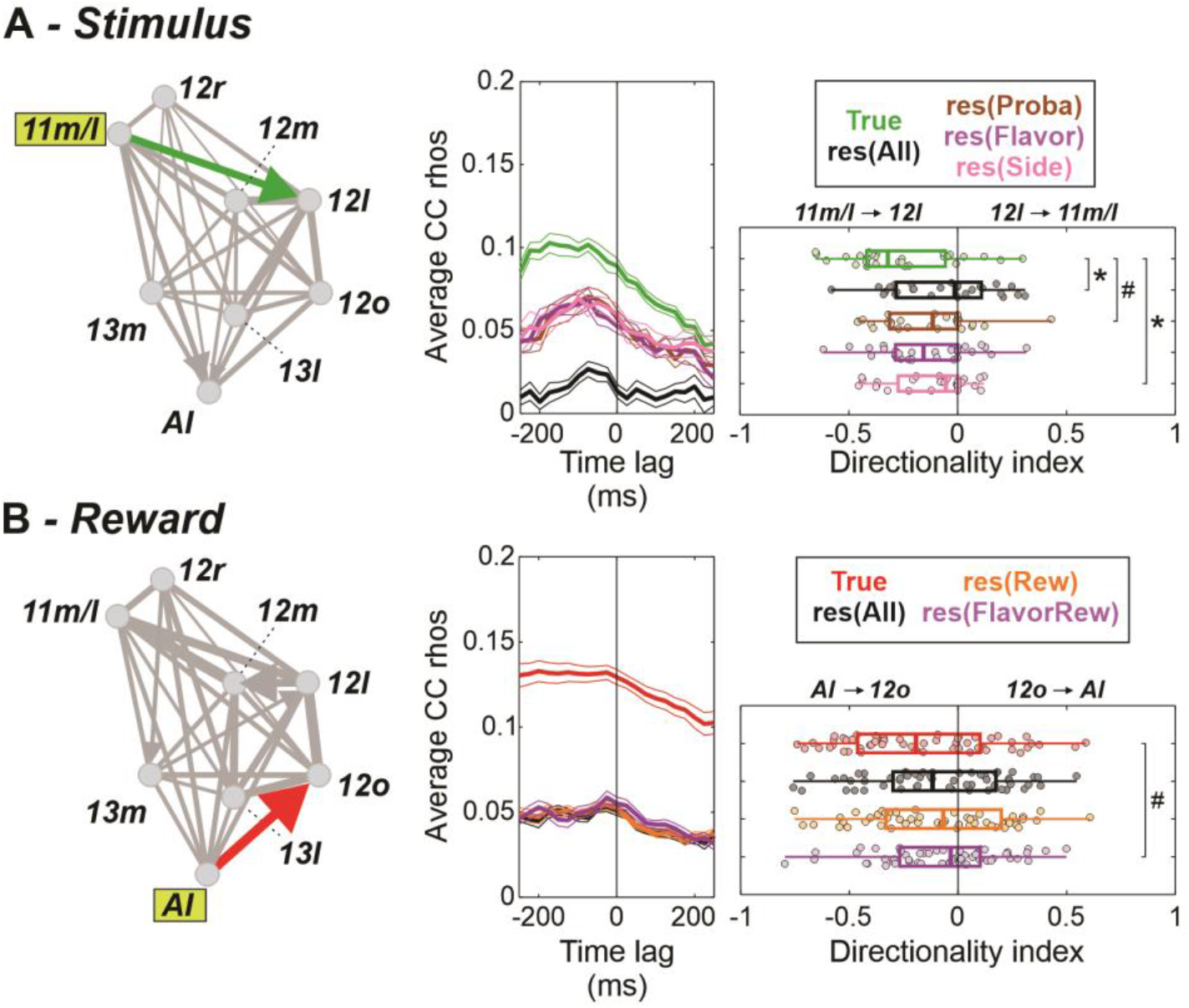
Connectivity lags are not entirely related to decision variable representations. Related to Figure 6. (**A**) Graph representation of the average correlation during the stimulus period and highlighting the 11m/l to 12l lag in connectivity, adapted from **Figure 6A**. Middle panel show the average correlation across different time lags when using the full neural activity (green) or when using the residual neuronal activities from 4 different models. Models used to extract residual activity included the chosen probability (brown), the chosen flavor (violet), the chosen side (pink), all three variables (black) or neither (green, as in **Figure 5**). These models were only applied to the activity of neurons in area 11m/l (yellow box in left panel). Right panel show the distribution of directionality indices across sessions (dots) and conditions (using 11m/l residual activity or not). Stars in right panels indicate a statistical bias in the directionality index across sessions when comparing the directionality in connectivity observed when using residuals compared to the true connectivity (Wilcoxon signed rank test; # p<0.1, * p<0.05, ** p<0.01 and *** p<0.001). (**B**) As panel A but for the reward period, where we focused on the connectivity between AI and area 12o. In this case, models used to extract residual activity in AI (yellow box on left panel) included whether monkeys were rewarded (orange), the received reward flavor when reward was delivered (violet), both variable with received flavor nested under reward (black) or neither (red, as in **Figure 5**).

**Table S1.** Number of recorded neurons for each monkey and area. Related to Figure 1.

| Area | vIPFC |  |  |  | OFC |  |  | AI |
| --- | --- | --- | --- | --- | --- | --- | --- | --- |
|  | 12r | 12m | 12o | 12l | 11m/l | 13l | 13m |  |
| mk M | 241 | 129 | 260 | 278 | 820 | 297 | 373 | 347 |
| mk X | 195 | 292 | 885 | 497 | 281 | 514 | 337 | 538 |

**Table S2.** Number of sessions included in the CCA analyses. Related to Figure 2. Numbers are indicated for each monkey separately (M-X). **#** indicates the pair of areas not considered for further analyses as not enough sessions were available.

| Number of sessions (M-X) | Area #2 |  |  |  |  |  |  |  |  |
| --- | --- | --- | --- | --- | --- | --- | --- | --- | --- |
| Area #1 |  | 12r | 12m | 12o | 12l | 11m/l | 13l | 13m | AI |
|  | 12r | . | . | . | . | . | . | . | . |
|  | 12m | 1-0 (#) | . | . | . | . | . | . | . |
|  | 12o | 5-0 | 0-13 | . | . | . | . | . | . |
|  | 12l | 8-0 | 0-4 | 3-16 | . | . | . | . | . |
|  | 11m/l | 18-0 | 1-3 | 19-19 | 23-5 | . | . | . | . |
|  | 13l | 5-0 | 1-9 | 5-42 | 6-3 | 19-13 | . | . | . |
|  | 13m | 9-0 | 1-4 | 10-26 | 9-2 | 40-11 | 9-16 | . | . |
|  | AI | 9-0 | 0-6 | 11-44 | 8-2 | 33-10 | 9-31 | 17-18 | . |

